# Cryo-EM structures reveal the mechanism of phosphatidylserine remodeling by membrane-bound glycerophospholipid *O*-acyltransferase 1

**DOI:** 10.64898/2026.09.01.748671

**Authors:** Leehyeon Kim, Siyoung Kim, Ritchie Ly, Luke Cohen-Abeles, Pei Liu, Xuejun Jiang, Dohoon Kwon, Robert V. Farese, Tobias C. Walther

## Abstract

Lands cycle remodeling of glycerophospholipid acyl chains is crucial for cells to maintain appropriate membrane composition. Glycerophospholipids are cleaved at the glycerol *sn*2-position by phospholipase A. The lysophospholipids are reacylated by enzymes of the membrane-bound *O*-acyltransferase (MBOAT) family to incorporate specific fatty-acyl chains to adjust membrane properties. How MBOAT enzymes recognize specific acyl-CoA donors, select lysophospholipid acceptors, and release products is unclear. Phosphatidylserine (PS), a critical anionic phospholipid, controls membrane surface charge, signaling-protein recruitment, and cell-death-associated membrane recognition, and PS acyl-chain remodeling is linked to ferroptosis resistance. Here, we showed that MBOAT1 preferentially generates monounsaturated fatty acid– containing PS from lyso-PS. High-resolution cryo-electron microscopy structures of human MBOAT1 captured distinct binding poses of the fatty acyl donor, lyso-PS acceptor, and PS product. With lipidomics, enzymology and molecular dynamics simulations, these structures reveal the mechanism and pathway of MBOAT1-dependent PS remodeling.

## Introduction

Acyl chains esterified to glycerophospholipids are critical determinants of membrane physical properties. While phospholipid head groups are initially determined during *de novo* glycerophospholipid synthesis, their acyl chains are continuously remodeled through a series of reactions known as the Lands’ cycle^1–3^. In this pathway, phospholipase A generates lysophospholipids that are subsequently reacylated with a specific fatty acyl chain. Such remodeling generates diverse phospholipid species and allows cells to adjust membrane composition according to biophysical and physiological demands. For instance, selective incorporation of polyunsaturated fatty acids into phospholipids regulates membrane fluidity and modulates cellular sensitivity to ferroptosis, an iron-dependent form of regulated cell death driven by lipid peroxidation^4^. Therefore, the enzymes that catalyze reacylation of lysophospholipids and their specificity for different substrates are essential for membrane homeostasis. However, the molecular mechanisms governing their selectivity remain poorly understood.

Reacylation of lysophospholipids is catalyzed by members of the membrane-bound *O*-acyltransferase (MBOAT) superfamily of enzymes^5–7^. These integral membrane proteins transfer fatty acyl groups from acyl-CoA donors to lipid or protein acceptors. MBOAT family members, including ACAT1, DGAT1, HHAT, PORCN, MBOAT5, and MBOAT7, have a shared transmembrane scaffold with an evolutionarily conserved catalytic histidine and provided important insights into the mechanism of acyl transfer^8–16^. However, despite recent advances in the structural understanding of MBOAT family enzymes, it is not understood how enzymes sharing a conserved MBOAT fold achieve distinct substrate specificities, distinguish lysophospholipid head groups, select specific acyl-CoA donors, and coordinate substrate entry with product release within the membrane.

MBOAT1 has been implicated in oleoyl-CoA-dependent phospholipid remodeling (Fig.1a). Previous studies suggest that MBOAT1 deficiency alters phosphatidylserine (PS) and phosphatidylethanolamine (PE) species^6^, in agreement with a role in monounsaturated fatty acid incorporation into glycerophospholipids. This distinction is important because PS and PE have fundamentally different cellular functions. PS is a quantitatively minor anionic phospholipid that regulates membrane surface charge, signaling-protein recruitment, and cell-death-associated membrane recognition^17,18^. PE is a major structural phospholipid involved in membrane curvature and bilayer organization^19^. Thus, defining whether MBOAT1 directly remodels PS, PE, or both is essential for understanding the physiological function of this enzyme. By replacing oxidation-prone polyunsaturated acyl chains with monounsaturated acyl chains, MBOAT1-dependent membrane remodeling dilutes the pool of lipids susceptible to ferroptotic lipid peroxidation^4^. However, cellular lipidomes reflect the integrated output of many reactions, complicating the interpretation of effects caused by individual lipid enzymes. It therefore remains unknown which lyso-phospholipid species are directly recognized by MBOAT1 as its physiological substrate, how it selects oleoyl-CoA as its preferred acyl donor, and how substrate entry and product release are coordinated in the membrane environment.

**Fig. 1.**
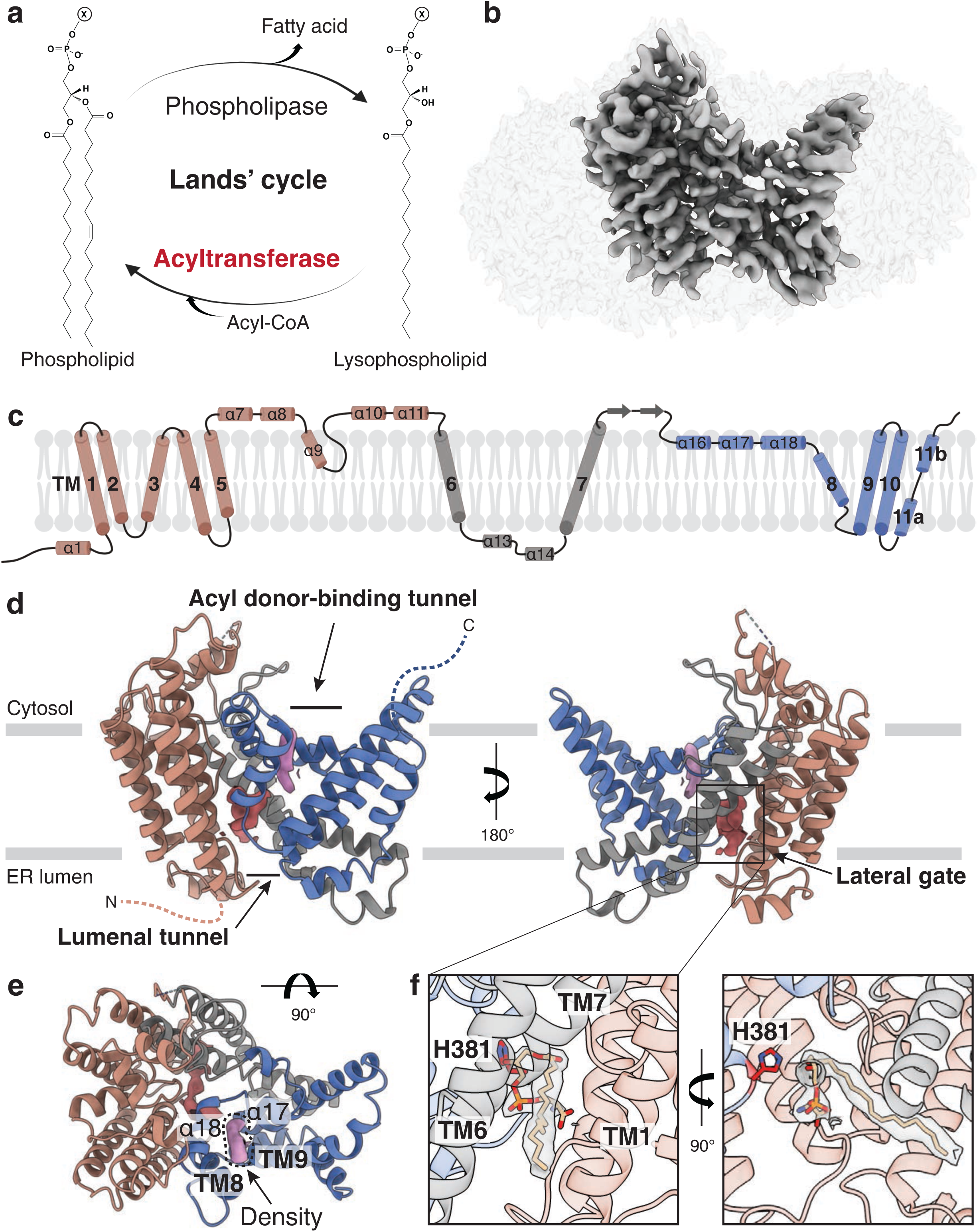
Overall structure of human MBOAT1. **a,** Schematic of the Lands’ cycle. Phospholipases remove a fatty acyl chain from phospholipids to generate lysophospholipids, which are reacylated by acyltransferases using acyl-CoA donors. **b,** Cryo-EM density map of human MBOAT1. **c,** Membrane topology of human MBOAT1. Transmembrane helices are numbered and colored according to the structural model, with the N-terminal lobe, central connecting region and C-terminal lobe shown in pink, grey and blue, respectively. **d,** Overall structure of human MBOAT1 shown in two orthogonal views within the ER membrane, with the acyl donor-binding tunnel, lateral gate, and lumenal tunnel indicated. MBOAT1 adopts a membrane-embedded MBOAT fold composed of an N-terminal lobe, a central connecting region and a C-terminal lobe. Non-proteinaceous densities observed in the wild-type map are shown as surfaces in red and purple. **e,** Close-up top view of the endogenous lipid-like density observed near the acyl donor-binding tunnel. The density is surrounded by α17, α18, TM8 and TM9. **f,** Local refinement improves the density for the co-purified lysophospholipid-like molecule. The ligand near the catalytic His381 is accommodated in a lateral gate formed by TM1, TM6 and TM7. Two orthogonal views are shown, related by a 90° rotation.

Here, we address these questions by combining cryo-electron microscopy (cryo-EM), quantitative biochemical analyses, molecular dynamics (MD) simulations, and cell-based lipidomics. Our findings reveal the molecular mechanisms of MBOAT1 substrate selectivity and more broadly provide a detailed mechanistic framework for lipid remodeling by MBOAT enzymes.

## Results

### A cryo-EM structure of human MBOAT1

We expressed human MBOAT1 in HEK293S GnTI^-c^ells and isolated the detergent-solubilized protein by affinity and size-exclusion chromatography. The purified protein eluted as a monodisperse species at a time consistent with the monomeric size of MBOAT1 (Extended Fig. 1a,b).

Using these protein preparations, we determined a cryo-EM map of human MBOAT1 at 3.07Å resolution. This map enabled us to build *de novo* a nearly complete model comprising residues 13–471 (Fig. 1b, Extended Data Figs. 2-4 and Table 1). MBOAT1 contains 11 transmembrane segments, including an N-terminal lobe, a central connecting region, and a C-terminal lobe that come together in the evolutionarily conserved MBOAT fold, situating the enzyme within the membrane (Fig. 1c). The overall architecture of wild-type (WT) MBOAT1 is similar to previously determined MBOAT family members^15,16^. The structure reveals the canonical access features of the MBOAT fold, including a cytoplasm-facing acyl donor-binding tunnel, a membrane-embedded lateral gate, and a lumenal tunnel that opens toward the ER lumen (Fig. 1d).

Analysis of the cryo-EM map of WT MBOAT1 revealed two non-proteinaceous densities (Fig. 1d). The first, less well-defined density was surrounded by α17, α18, TM8 and TM9, corresponding to the putative acyl donor-binding site. However, the map quality was insufficient for unambiguous ligand assignment or model building (Fig. 1e).

A second density was located in close proximity to the catalytic His381 and likely represents an endogenous lipid co-purified with MBOAT1 (Fig. 1d). Focused refinement improved the local density around this putative ligand, revealing an elongated molecule accommodated in a lateral gate formed by transmembrane helices 1, 6, and 7 (Fig. 1f), lined predominantly by hydrophobic and aromatic residues (e.g., Ile29, Phe36, Val37, Trp136, Met143, Phe255, Lys295, Tyr296, Phe298, Ala299, Trp300, Trp380 and Phe441) (Extended Data Fig.1c). The dimensions and geometry of the density are consistent with a lysophospholipid, and its protein environment included a defined head-group-binding region and an acyl chain-accommodating hydrophobic tunnel extending toward the membrane bilayer. Mass spectrometric (MS) analysis of co-purified lipids detected a signal most consistent with 18:0 lyso-PS in preparations of MBOAT1, but not in similarly prepared MBOAT2 or MBOAT5 samples (Extended Data Fig. 1d,e). MS/MS fragmentation of this ion matched reference spectra of lyso-PS.

### MBOAT1 is primarily a lyso-phosphatidylserine acyltransferase

To test whether MBOAT1 activity is specific for PS remodeling, we measured the effect of MBOAT1 deletion on the cellular lipidome of T47D breast cancer cells where MBOAT1 has been linked to ferroptosis surveillance^4^. Cells lacking MBOAT1 specifically had lower levels of MUFA-containing PS species, including PS 18:0/18:1 and PS 16:0/18:1, than WT cells (Fig. 2a). Consistently, analysis of the relative acyl-chain distribution within the total PS pool revealed a shift forward PUFA-containing species in MBOAT1-deficient cells (Fig. 2b)

**Fig. 2.**
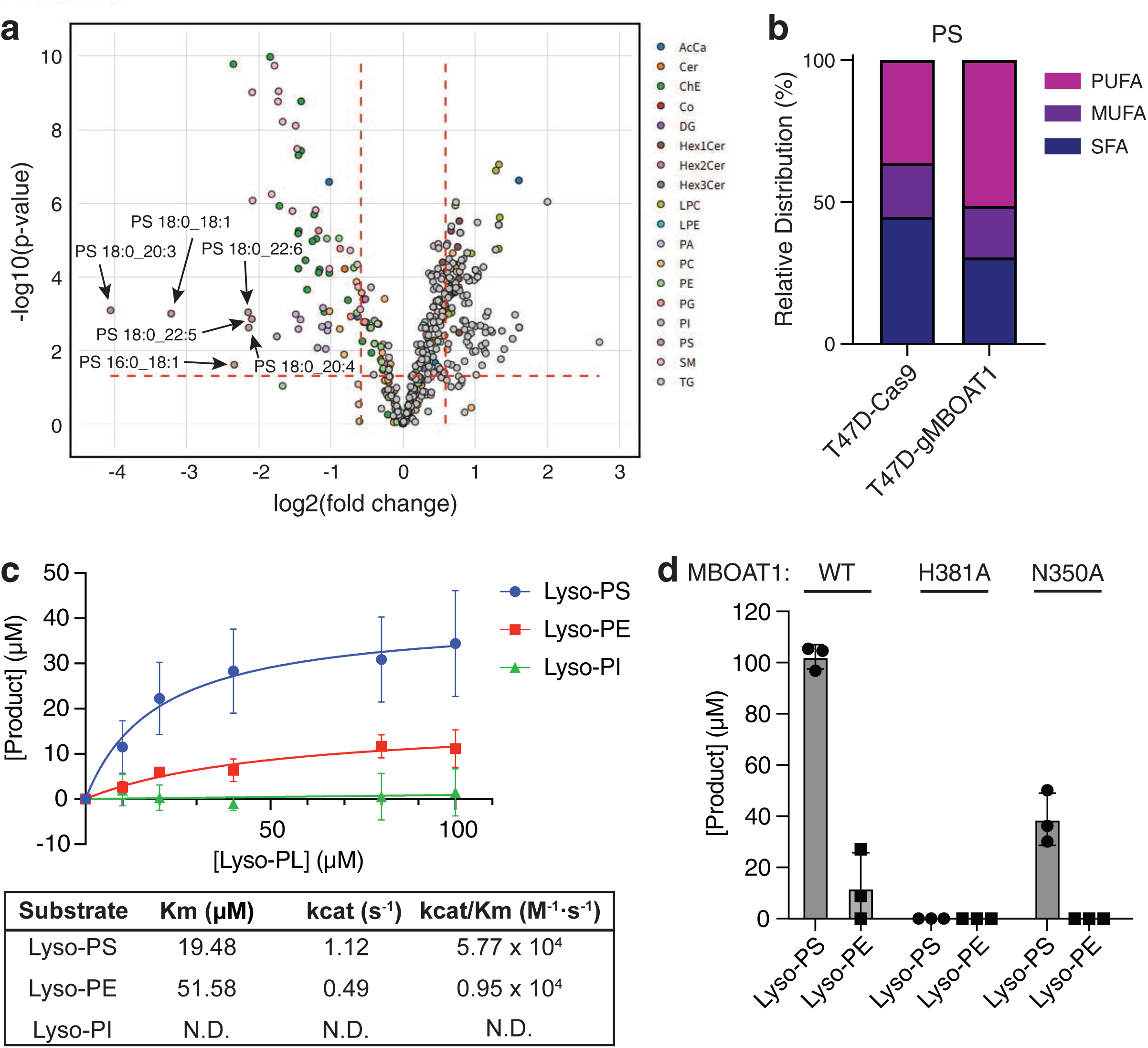
MBOAT1 preferentially acylates lyso-PS in cells and *in vitro*. **a,** Volcano plot of lipidomic changes in MBOAT1-deficient T47D cells compared with WT cells. Selected PS species reduced upon MBOAT1 loss are labelled. Dashed lines indicate fold-change and significance thresholds. Cutoff: FC threshold = 1.5, p < 0.05; Welch’s t-test. **b,** Relative distribution of SFA, MUFA, and PUFA species within PS. **c,** Acyltransferase activity of purified MBOAT1 with lyso-PS, lyso-PE or lyso-PI as acyl acceptors and oleoyl-CoA as the acyl donor. Michaelis constant (Km) and turnover number (kcat) values for MBOAT1 with the indicated lysophospholipid substrates. Activities were measured with purified MBOAT1, 100 μM oleoyl-CoA and increasing concentrations of each lysophospholipid. N.D., not determined. **d,** Endpoint activity assay of purified MBOAT1 WT and catalytically impaired variants. Reactions contained 100 μM oleoyl-CoA and 100 μM lyso-PS or lyso-PE. Data are mean ± standard deviation (SD), n = 3 independent experiments.

In agreement with these data, we found high activity of purified MBOAT1 towards lyso-PS *in vitro.* Acyltransferase activity of purified MBOAT1 using different lyso-phospholipid acceptors and oleoyl-CoA as the acyl donor showed robust, saturable activity toward lyso-PS (kcat/Km = 5.77 x 10^4^ M^-1^ s^-1)^, but markedly lower activity for lyso-PE, with a higher apparent Km and lower kcat (kcat/Km = 0.95 x 10^4^ M^-1s-1;^ Fig. 2d). We also did not find activity of purified MBOAT1 when we provided lyso-PI as a substrate.

To independently validate these kinetic results, we directly quantified phospholipid product formation in endpoint assays. In agreement with the kinetic analysis, MBOAT1 robustly generated PS, whereas PE formation was minimal (Fig. 2e). In contrast, alanine mutants of the putative catalytic residues H381 and N350 showed little to no detectable product formation (Fig. 2d, Extended Data Fig.1a,b).

### Lysophospholipids enters MBOAT1 through a lateral gate

We next sought to define the entry route of lyso-PS into the deeply buried active site of MBOAT1 and how the acyl acceptor is positioned for catalysis. For most lipid-modifying MBOAT enzymes, two possibilities may explain how acyl-acceptor substrates reach the catalytic center: either through an ER lumen-facing tunnel or a membrane bilayer-facing lateral gate (Fig. 1d). To capture intermediate states along the lyso-PS entry pathway, we determined cryo-EM structures of catalytically inactive MBOAT1 mutants supplemented with 16:0 lyso-PS immediately before vitrification. Specifically, we analyzed H381A and N350A mutants, in which the catalytic histidine or asparagine, respectively, was replaced by alanine (Fig. 3a,b).

**Fig. 3.**
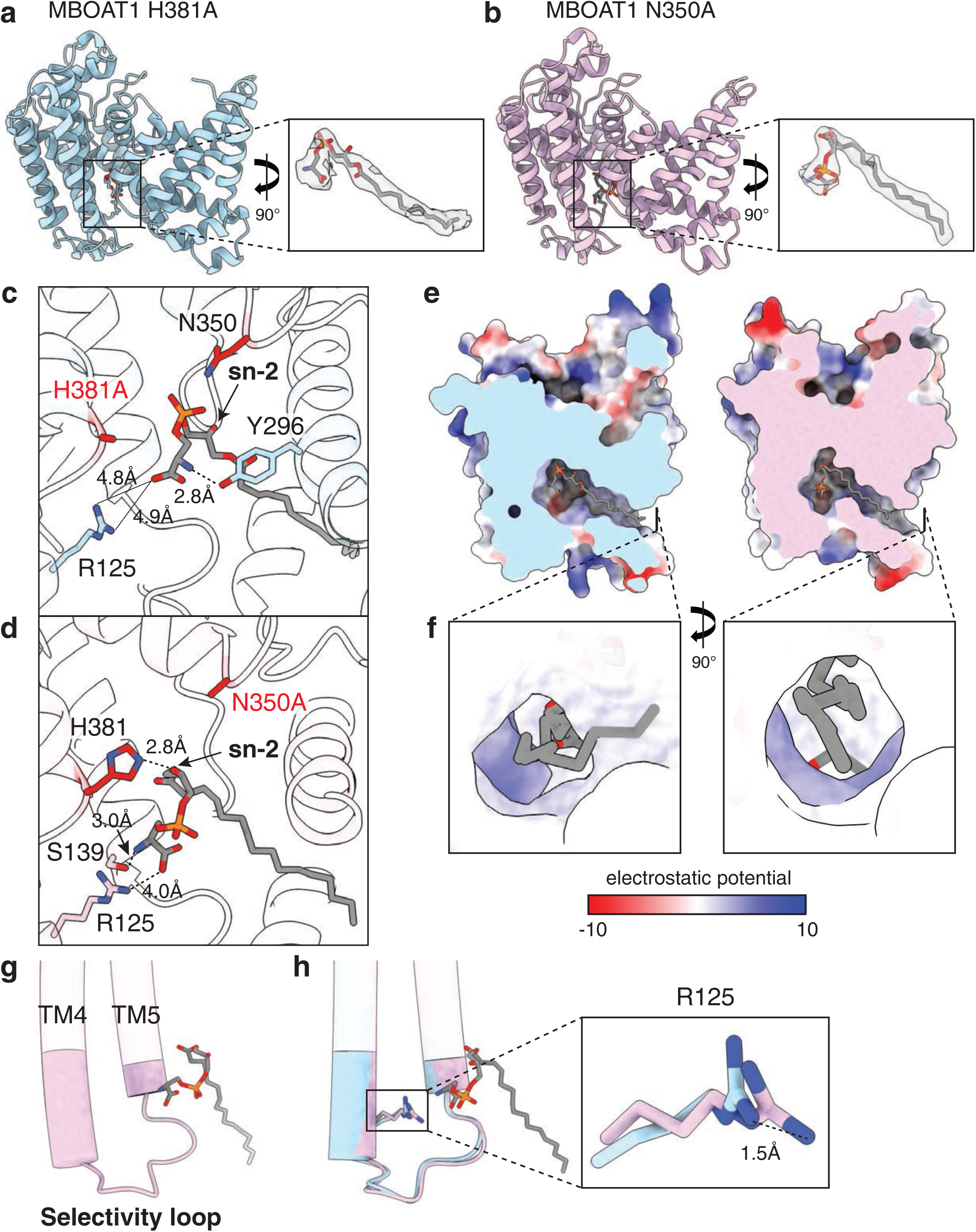
Lyso-PS entry and catalytic positioning in MBOAT1. **a,** Cryo-EM structure of the MBOAT1 H381A mutant bound to lyso-PS, capturing a partially inserted entry state. Insets show the lyso-PS density and model along the lateral entry pathway in two orthogonal views. **b,** Cryo-EM structure of the MBOAT1 N350A mutant bound to lyso-PS, capturing a fully engaged acceptor-bound state. Insets show the lyso-PS density and model near the catalytic center in two orthogonal views. **c,** Close-up view of lyso-PS in the H381A structure. Lyso-PS has entered the internal cavity but is displaced from the catalytic center, with the acyl chain only partially accommodated in the hydrophobic tunnel and the glycerol backbone offset from the productive pose. **d,** Close-up view of lyso-PS in the N350A structure. The acyl chain extends into the hydrophobic tunnel, and the glycerol *sn*-2 hydroxyl is positioned adjacent to His381 and Asn350, consistent with a catalytically competent acceptor pose. **e,** Electrostatic surface representation of the lateral entry pathway shown in two orthogonal views. The membrane-facing entrance contains an electropositive region that leads into a hydrophobic tunnel toward the catalytic cavity. Electrostatic potential is colored from red (negative) to blue (positive) as indicated. **f,** Close-up view of the lateral entry region in the N350A and H381A structures, highlighting distinct lyso-PS positions along the entry pathway. **g,** Location of the selectivity loop between TM4 and TM5, comprising residues 120–140 in MBOAT1. **h,** Arg125 side-chain shift between the H381A and N350A structures. In the N350A acceptor-bound state, Arg125 moves approximately 1.5Å closer to the lyso-PS head group.

The cryo-EM structure of MBOAT1 H381A captured lyso-PS in a partially inserted pose along the lateral gate entry pathway (Fig. 3a). The phosphoserine head group of lyso-PS had entered the lateral gate and was positioned to interact with Try296 and Arg125, but the substrate did not advance into the fully engaged catalytic pose. The acyl chain was only partially accommodated within the hydrophobic gate, and the glycerol backbone remained displaced from the catalytic center (Fig. 3c). Consequently, the *sn*-2 hydroxyl group was not positioned for productive acyl transfer, and we therefore interpret it as a mutant-trapped, nonproductive substrate-entry state.

Compared with the MBOAT1 H381A entry state, the cryo-EM structure of MBOAT1 N350A captured a more deeply engaged lyso-PS pose (Fig. 3b). The non-proteinaceous density was more clearly resolved than in the initial WT MBOAT1 structure and could be modelled well as a lyso-PS molecule. As lyso-PS advanced into the acceptor-binding site, the phosphoserine head group moved closer to Arg125 and Ser139. The lyso-PS acyl chain extended further into the protein through the membrane-facing lateral gate, positioning the glycerol *sn*-2 hydroxyl group adjacent to His381 (Fig. 3d). The position and geometry of the substrate were consistent with a catalytically competent acceptor pose, in which His381 was positioned to activate the lyso-PS *sn*-2 hydroxyl group for nucleophilic attack on the acyl-CoA thioester.

These structures revealed a lateral gate as an entry pathway for lyso-PS and showed how the substrate advances from a partially inserted entry state to a catalytically competent acceptor pose. In this model, the lateral opening between TM1 and TM6 connects the membrane bilayer-facing surface of MBOAT1 to its intramembrane catalytic center. Residues Leu27, Gln33, Phe36, Val37, Leu252, Phe255, Lys295 and Ala299 define the walls of the lateral gate, forming a continuous passage from the lumenal leaflet of the ER membrane to the catalytic center.

Electrostatic surface analysis of the lateral gate revealed an electropositive surface at the membrane-facing entrance to the hydrophobic tunnel (Fig. 3e). This positively charged environment may facilitate entry of the anionic phosphoserine head group towards the catalytic center, while the hydrophobic tunnel accommodates the acyl chain (Fig. 3f).

Adjacent to the phosphoserine head-group binding site, MBOAT1 contains a loop connecting TM4 and TM5 that was previously identified by loop-swapping studies as a determinant of phospholipid head group specificity^16^ (Fig. 3g, Extended Data Fig. 5). We refer to this region, comprising residues 120–140 in MBOAT1, as the selectivity loop. Comparison of the cryo-EM structures of MBOAT1 H381A and N350A revealed no major change in the overall conformation of the selectivity loop. In contrast, the side chain of Arg125 shifted by approximately 1.5 Å towards the phosphoserine head group as the substrate advanced into the acceptor-binding site (Fig. 3h). Although the distance is too great to support a stable salt bridge, the progressive repositioning of Arg125 suggests that it transiently guides the phosphoserine head group towards its catalytically competent position.

### Acyl donor tunnel geometry underlies oleoyl-CoA selectivity of MBOAT1

MBOAT1 has a strong preference for oleoyl-CoA as acyl donor substrate over other acyl-CoA species, including polyunsaturated arachidonoyl-CoA or short-chain acetyl-CoA^6,7^. To define the structural basis of this preference, we determined the cryo-EM structure of MBOAT1 N350A in the presence of oleoyl-CoA. This mutant retains lyso-PS and acyl-donor binding but prevents productive catalysis, thereby stabilizing a donor-bound state (Fig. 4a and Extended Data Figs. 2–4).

**Fig. 4.**
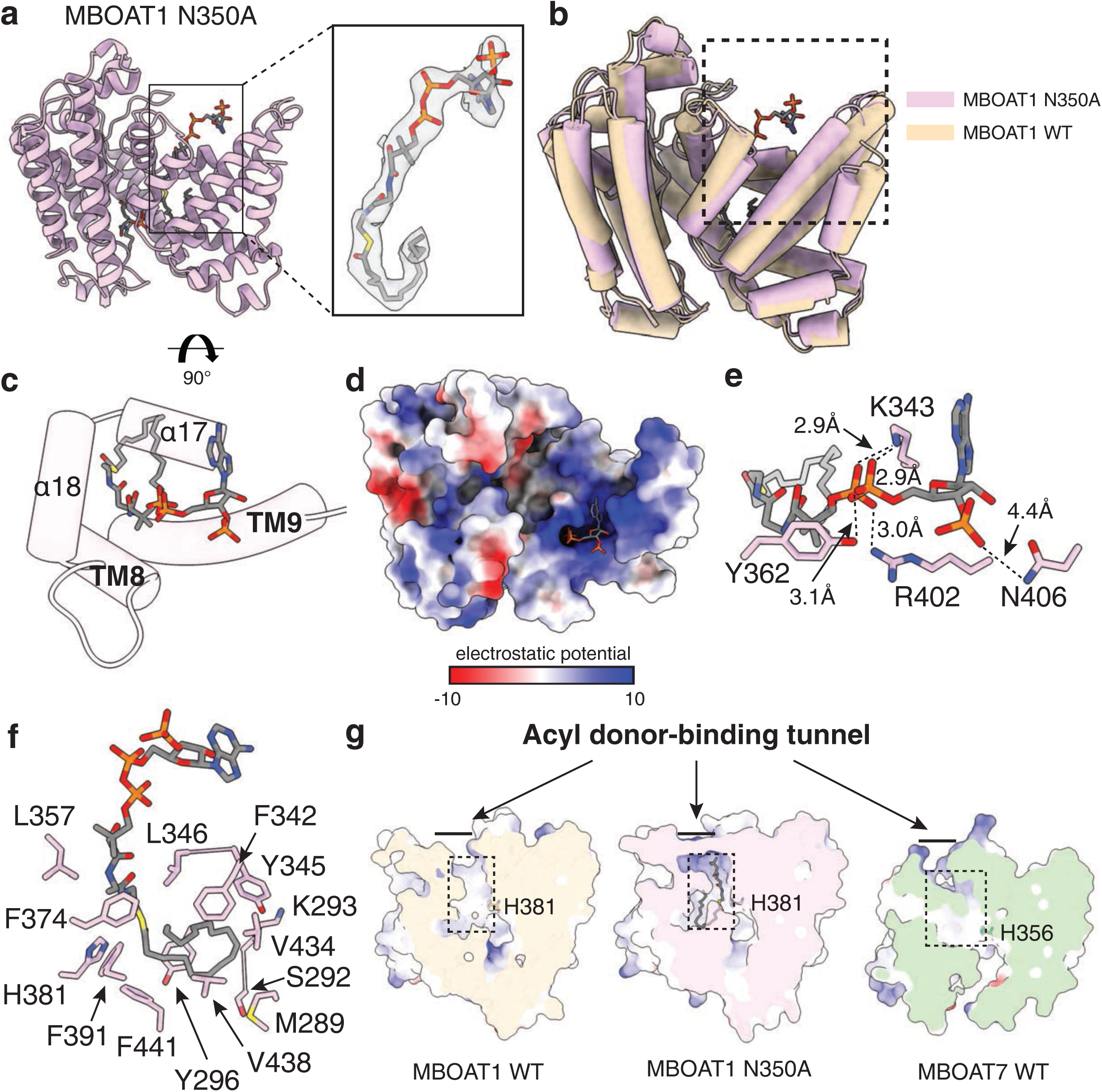
Structural basis for oleoyl-CoA recognition by MBOAT1. **a,** Cryo-EM structure of MBOAT1 N350A simultaneously bound to lyso-PS and oleoyl-CoA. MBOAT1 is shown as a cartoon, and lyso-PS and oleoyl-CoA are shown as sticks. **b,** Superposition of the MBOAT1 N350A complex and WT MBOAT1. The MBOAT1 N350A is shown in pink and WT MBOAT1 in pale yellow, highlighting the overall conservation of the transmembrane scaffold upon donor engagement. **c,** Cryo-EM density for oleoyl-CoA in the acyl donor-binding tunnel, shown in two orthogonal views. **d,** Electrostatic surface representation of the oleoyl-CoA-binding site. The CoA moiety is positioned at the cytoplasm-facing entrance of the acyl donor-binding tunnel, and the oleoyl chain extends into a membrane-embedded hydrophobic tunnel. **e,** Close-up view of the CoA head-group-binding site. Lys343 and Arg402 form electrostatic and hydrogen-bonding interactions with the phosphate groups, whereas Tyr362 and Asn406 contribute additional polar contacts. Dashed lines indicate selected distance**. f,** Residues lining the oleoyl-chain-binding tunnel. The acyl chain is surrounded by residues, including Met289, Ser292, Lys293, Tyr296, Phe342, Tyr345, Leu346, Leu357, Phe374, His381, Phe391, Val434, Val438 and Phe441, forming an extended acyl donor-binding tunnel within the membrane-embedded core of MBOAT1. **g,** Comparison of the acyl donor-binding tunnel in WT MBOAT1, oleoyl-CoA-bound MBOAT1 N350A and WT MBOAT7 (PDB: 8ERC). The acyl-chain tunnel differs in shape and dimensions between MBOAT1 and MBOAT7, suggesting a structural basis for enzyme-specific acyl-CoA donor preference. Electrostatic potential is colored from red (negative) to blue (positive) as indicated.

Comparison of the cryo-EM structures of MBOAT1 WT and N350A revealed a high degree of structural similarity, with a Cα root-mean-square deviation (RMSD) of 1.22Å over the transmembrane core, indicating that donor binding does not require large-scale rearrangements of the MBOAT1 scaffold (Fig. 4b). Instead, oleoyl-CoA binding was accompanied by local tightening of the acyl donor binding tunnel. The cryo-EM map of MBOAT1 N350A revealed a continuous, elongated density that could be confidently modelled as oleoyl-CoA (Fig. 4c). This density corresponds to the acyl donor-binding cavity identified in the WT structure, which is formed by the C-terminal lobe, including α17, α18, TM8, and TM9 (Figs. 1e, 4c). The N350A structure, therefore, confirms this C-lobe cavity as the acyl donor-binding site of MBOAT1. Within this site, the CoA moiety is positioned at the cytoplasm-facing entrance, whereas the oleoyl chain extends deeply into the membrane-embedded core of the enzyme.

The CoA moiety is anchored at the entrance of the acyl donor binding tunnel by a cluster of polar and basic residues (Fig. 4d). The 3′-phosphoadenosine diphosphates moiety of CoA is coordinated primarily by Lys343 and Arg402, which form extensive hydrogen-bonding and electrostatic interactions with the phosphate groups (Fig. 4e). Tyr362 contacts the ribose-phosphate region, whereas Arg405 is positioned adjacent to the distal nucleotide moiety and contributes to the electropositive environment at the pocket entrance. Asn406 further contributes to the polar environment surrounding the CoA phosphate group. These interactions stabilize the CoA head group at the cytoplasm-facing entrance and orient the pantetheine arm into a narrow channel leading toward the catalytic center. Near the thioester linkage, the pantetheine arm passes adjacent to Trp349, positioning the thioester next to His381 and the Asn350 (Extended Data Fig. 6a).

The 18-carbon acyl chain with the characteristic single *cis* double bond at the Δ9 position extended deeply into the membrane-embedded core of the enzyme (Extended Data Fig. 6b). The acyl chain of oleoyl-CoA was accommodated within a predominantly hydrophobic tunnel lined by Met289, Ser292, Lys293, Tyr296, Phe342, Tyr345, Leu346, Leu357, Phe374, His381, Phe391, Val434, Val438 and Phe441 from TM7–TM10 (Fig. 4f). These residues form a continuous hydrophobic tunnel that accommodates the 18-carbon monounsaturated acyl chain. The tunnel follows the length and curvature of the oleoyl chain, featuring a local constriction that complements the bend imposed by the Δ9 cis double bond.

To understand how these structural features contribute to acyl-donor selectivity, we compared the acyl-donor binding tunnels of MBOAT1 and MBOAT7, the latter of which preferentially uses polyunsaturated acyl-CoA donors^16^. The acyl-chain tunnels of MBOAT1 and MBOAT7 differ markedly in shape (Fig. 4g). Compared with MBOAT7, MBOAT1 contains bulkier hydrophobic residues, including Phe342, Val313, Val438 and Cys411, that line and narrow the donor acyl-chain tunnel (Extended Data Fig. 6c). These residues create a narrower pocket that is less compatible with accommodation of acyl-CoA donors carrying multiple cis double bonds. By contrast, the corresponding acyl-chain tunnel in MBOAT7 is wider and more curved, consistent with accommodation of a polyunsaturated arachidonyl chain.

To compare how different MBOAT enzymes accommodate the CoA moiety of the acyl-donor substrate, we superimposed the structures of MBOAT1 N350A, MBOAT7 and MBOAT5 (Extended Data Fig. 6d). Electrostatic surface analysis revealed electropositive regions at the corresponding cytoplasm-facing CoA-binding entrances (Extended Data Fig. 6e), suggesting that these enzymes share similar electropositive surfaces for initial recognition of the CoA head-group. Although these overall electrostatic features are conserved, the detailed orientation of the nucleotide moiety differs markedly among MBOAT family members. In MBOAT1, the adenine moiety is accommodated in an acyl-donor binding tunnel extending upward from the diphosphate group (Fig. 4d). In contrast, in the arachidonyl-CoA-bound MBOAT5 structure, the adenine moiety adopts the opposite orientation (Extended Data Fig. 6e). The predicted CoA-binding region of MBOAT7 also does not align well with the MBOAT1 binding pose, suggesting that individual MBOAT enzymes recognize the CoA head group through similar electropositive surfaces while accommodating CoA in distinct binding orientations.

### Product-bound MBOAT1 reveals a lateral PS release pathway

After acyl transfer, the newly synthesized PS product must be released from the enzyme. To capture the post-catalytic state, we determined the cryo-EM structure of WT MBOAT1 incubated with both lyso-PS and oleoyl-CoA for 30 min to enrich the population of product-bound molecules before vitrification. This yielded a 3.42Å cryo-EM map in which an additional density corresponding to a PS molecule was resolved within the catalytic cavity (Fig. 5a).

**Fig. 5.**
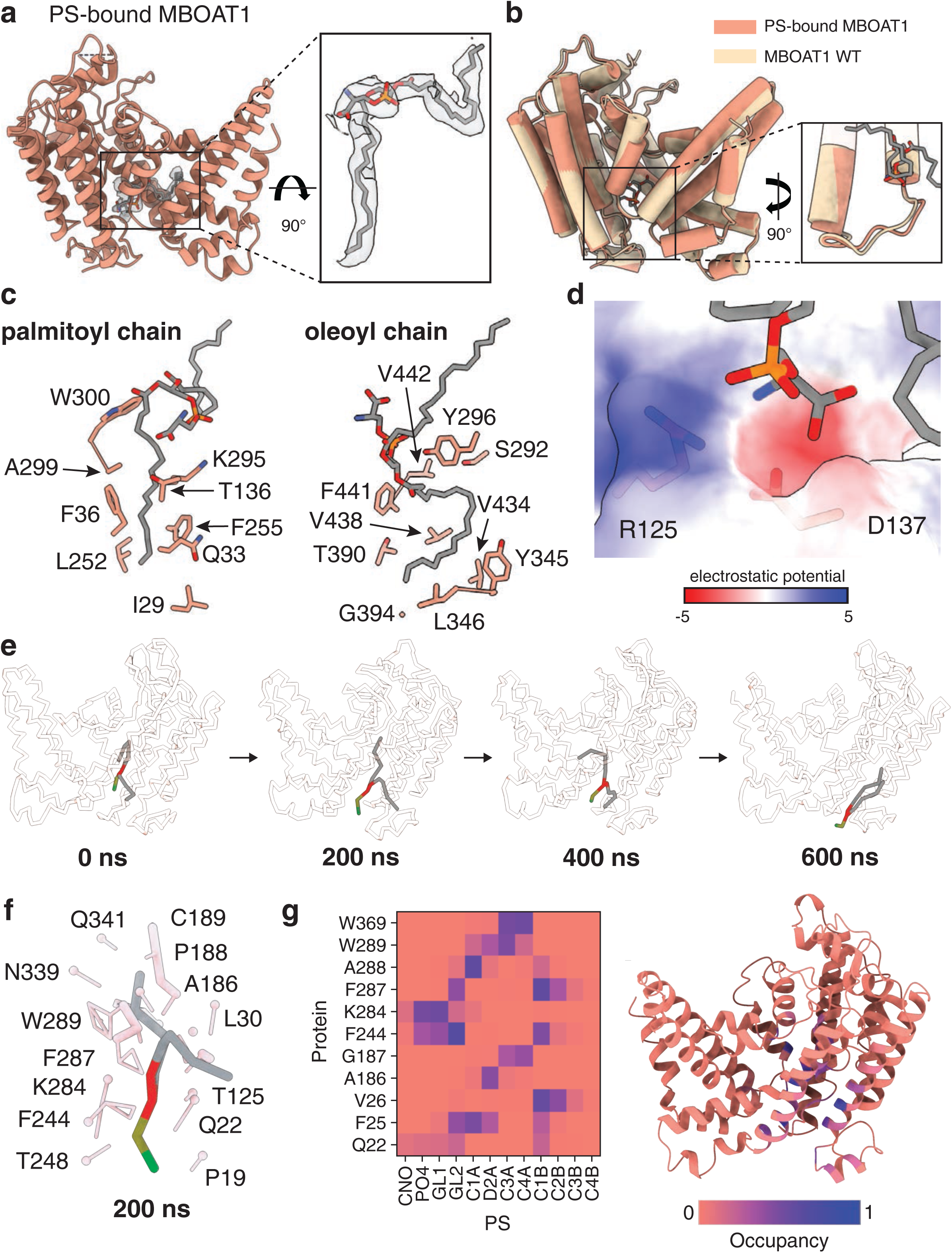
PS-bound MBOAT1 structure and product-release pathway. **a,** Overall structure of PS-bound MBOAT1, with PS shown as grey sticks. The PS density is shown as a grey mesh**. b,** Superposition of the PS-bound MBOAT1 complex and WT MBOAT1. The PS-bound MBOAT1 is shown in orange and WT MBOAT1 in pale yellow, highlighting the selectivity loop movement. **c,** Binding environment of the PS acyl chains. The 16:0 tail occupies the lateral gate, whereas the newly transferred 18:1 tail is accommodated in an adjacent acyl donor-binding tunnel. **d,** Electrostatic surface representation of the PS headgroup site. **e,** Representative molecular dynamics snapshots showing lateral movement of PS toward the membrane-facing exit pathway. **f,** Residues contacting PS in a representative 200-ns molecular dynamics snapshot. **g,** PS contact analysis from molecular dynamics simulations. Left, heat map of residue–PS contact occupancy. Right, contact occupancy mapped onto the MBOAT1 structure.

The overall conformation of product-bound MBOAT1 closely resembled that of the substrate-bound WT structure, with the transmembrane core largely unchanged (Fig. 5b; with a Cα RMSD of 0.88Å). By contrast, the selectivity loop showed a larger local rearrangement, with a Cα RMSD of 2.93Å. The product-bound structure contained a PS molecule within the catalytic cavity, with its two acyl chains extending into the hydrophobic interior of the enzyme (Fig. 5d). The acceptor-derived palmitoyl chain occupied the same tunnel engaged by the acyl chain of the lyso-PS substrate, whereas the acyl donor-derived oleoyl chain at the *sn*-2 position was accommodated in an adjacent hydrophobic groove that was vacant in the lyso-PS-bound state. This arrangement revealed how MBOAT1 accommodates the newly synthesized diacylated PS product while maintaining the phosphoserine head group in its recognition site. After acyl transfer, the negatively charged PS head group remained adjacent to Asp137 within the selectivity loop region (Fig. 5d). This configuration suggests that the product-bound state is poised for PS release rather than stable product retention. In agreement with a functional role for Asp137, a D137A mutation markedly reduced MBOAT1 activity *in vitro* (data not shown).

Since MBOAT1 contains both a lateral gate and a lumenal tunnel (Fig. 1d), the route by which the PS product exits the catalytic cavity was unclear. We, therefore, performed molecular dynamics (MD) simulations using the product-bound MBOAT1 structure. In coarse-grained (CG) MD simulations, the PS product was released into the membrane within 800 ns through the lateral gate (Fig. 5e) and ultimately partitioned into the lumenal membrane leaflet (Extended Movie 1). During this process, PS transiently adopted an intermediate conformation, as captured in the 200-and 400-ns snapshots (Fig. 5e). In this intermediate conformation, PS was stabilized by an electrostatic interaction with Lys284 and hydrophilic interactions with Thr248, Gln22, and Thr125, and its tails were surrounded by hydrophobic residues (Fig. 5f). The selectivity loop remained adjacent to this intermediate, suggesting that it contributes to guiding PS release. Before complete release of the PS into the membrane, PS bound non-specifically between helices 1 and 6, as shown in the 600-ns snapshot (Fig. 5e). To further characterize PS-protein interactions along the release pathway, we calculated an occupancy map. In agreement with the observed release trajectory, the occupancy map revealed persistent contacts between the PS and Lys284, and the acyl chains primarily interacted with hydrophobic residues lining the lateral gate (Fig. 5g).

## Discussion

Here, we report a series of cryo-EM structures of human MBOAT1 in complex with its substrates and products. Collectively, these structures define the mechanisms of substrate selectivity, catalysis and product release for this enzyme that generates PS.

Previously, lipidomic analyses of MBOAT1-deficient cells^4^ and enzymatic studies with microsomal fractions suggested the enzyme remodels both PS and PE species^6^. However, these studies could not determine whether MBOAT1 directly acts on lyso-PS, lyso-PE or multiple lysophospholipid substrates. We found that purified MBOAT1 has a strong intrinsic preference for lyso-PS, with only low activity toward lyso-PE *in vitro*. Consistently in breast cancer cells where MBOAT1 is important for ferroptosis protection, its deficiency affects almost exclusively PS acyl chain composition. Nevertheless, with more PE than PS in cellular membranes^20^, a minor contribution of MBOAT1 to PE remodeling *in vivo* cannot be excluded. The relevant substrate pool, however, is the local ER lyso-phospholipid pool, and quantitative information on ER lyso-PS and lyso-PE abundance is limited. Future ER-specific lipidomic analyses will be important for defining the physiological substrates available to MBOAT1.

With the different cryo-EM structures, a model for MBOAT1-mediated catalysis emerged (Fig. 6). Initially, MBOAT1 captures lyso-PS at the lateral-gate entrance into the enzyme, where the phosphoserine head group of the lipid engages with the selectivity loop of the protein (represented by the cryo-EM structure of MBOAT1 H381A). In a next step (represented in the cryo-EM structure of MBOAT1 N350A), lyso-PS advances into a catalytically competent pose, with its acyl chain fully accommodated in the hydrophobic tunnel and its *sn*-2 hydroxyl positioned near His381. Oleoyl-CoA binds from the cytoplasmic acyl-donor site, placing its thioester adjacent to the acceptor hydroxyl group, thereby aligning the two substrates for acyl transfer within the catalytic core of the membrane.

**Fig. 6.**
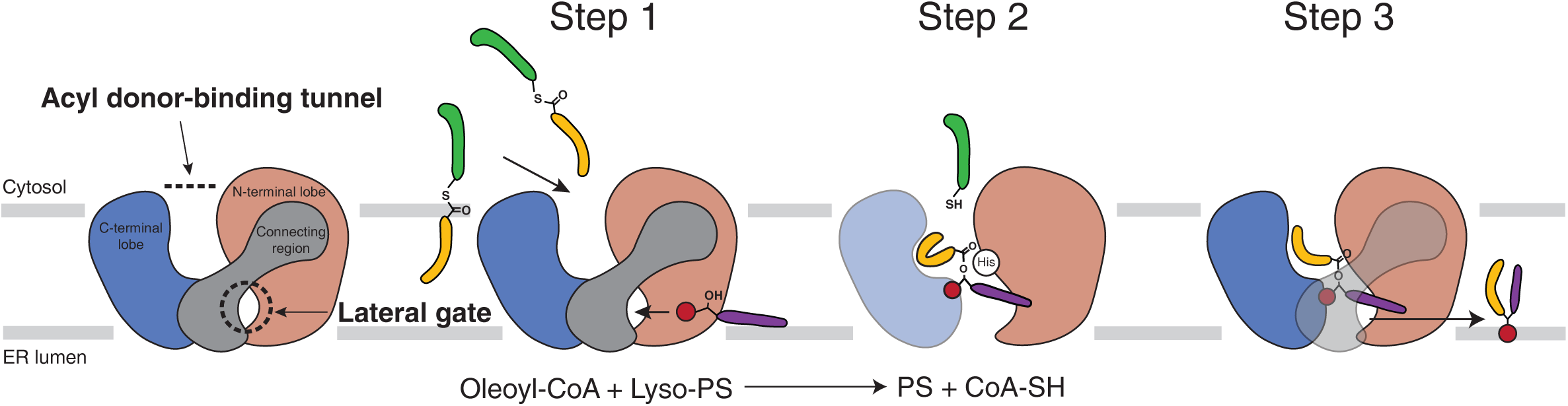
Structure-based working model for MBOAT1. The lateral gate and acyl donor-binding tunnel provide entry routes for lyso-PS and oleoyl-CoA, respectively. Lyso-PS and oleoyl-CoA bind at the membrane-embedded catalytic center (step 1). Acyl transfer is catalyzed by the conserved His381 residue, which promotes reaction of the lyso-PS *sn*-2 hydroxyl group with the oleoyl-CoA thioester (step 2). The PS product subsequently diffuses into the lumenal leaflet through the lateral gate (step 3).

After acyl transfer, the newly formed PS is accommodated in the internal cavity of MBOAT1, but its head group adopts an electrostatic suboptimal arrangement near Arg125 and Asp137 within the selectivity loop. In particular, the proximity of Asp137 to the negatively charged PS head group likely disfavors stable product retention. Thus, the same region that guides lyso-PS entry into the enzyme and positioning does not appear to stably retain the diacylated PS product after the reaction. This principle is reminiscent of the electrostatic product–release mechanism proposed for the PS synthesis enzyme PSS1, in which the reaction product is released from the catalytic cavity owing to repulsive electrostatic interactions with PS^21^. Together with MD simulations showing lateral movement of PS, these data suggest that MBOAT1 uses the selectivity loop region to couple substrate recognition, catalytic turnover and product release during phospholipid remodeling.

This catalytic mechanism for MBOAT1 action provides a framework for understanding substrate specificity across enzymes of the MBOAT family. Protein-acylating MBOATs, such as HHAT and PORCN, contain enlarged lumenal tunnels that likely accommodate protein substrates (Extended Data Fig. 7a), whereas the lateral gate used in MBOAT1 for lyso-PS entry is restricted or occluded (Extended Data Fig. 7b). DGAT1, which uses diacylglycerol as a lipid acceptor, shows a large lateral gate, consistent with entry of a bulky neutral lipid substrate (Extended Data Fig. 7c). In comparison, MBOAT1 contains a narrower lateral gate, shaped by TM1, TM2 and the selectivity loop region, that is suited for entry and positioning of a lysophospholipid acceptor. These comparisons suggest that MBOAT enzymes preserve a common catalytic scaffold while diversifying their substrate-access routes to match distinct acceptor chemistries.

Comparison with MBOAT5 and MBOAT7 further suggests how related lipid-remodeling MBOATs achieve distinct substrate and donor preferences. Although MBOAT1, MBOAT5 and MBOAT7 shared an overall fold, their lateral gates, selectivity loop regions and acyl-donor tunnels differ substantially. The lyso-PC-bound MBOAT5 structure had a more open lateral gate and a looser selectivity loop configuration than MBOAT1, whereas the corresponding region in MBOAT7 was displaced outward, leaving the acceptor-binding region more open (Extended Data Fig. 7d). For acyl donor binding, MBOAT5 and MBOAT7 contained broader, more curved acyl-chain tunnels than the constrained tunnel of MBOAT1 harboring oleoyl CoA, consistent with their use of polyunsaturated acyl-CoA donors. Thus, MBOAT acyl-chain specificity was likely encoded by local differences in lateral-gate architecture and acceptor-site positioning for the lyso-lipid substrate and acyl donor-tunnel geometry.

The topology of MBOAT1 within the ER raises questions about the membrane context of PS remodeling. Although most cellular PS resides in the cytoplasmic leaflet of the secretory pathway and plasma membrane^22^, recent structural work on human PSS1 suggested that it delivers newly synthesized PS initially to the lumenal leaflet of the ER membrane^21^. Because the MBOAT1 lateral gate is positioned near the lumenal side, MBOAT1 may remodel a lumenally accessible lyso-PS pool within the ER membrane. One possibility is that newly synthesized PS is locally deacylated in the lumenal leaflet to generate lyso-PS, which is then reacylated by MBOAT1. This would provide a local route for PS remodeling without requiring extensive interleaflet redistribution of PS before MBOAT1 engagement. Of note, PS 18:0/18:1 is thought to be an important component of the inner leaflet of cell membranes^23^. Likely, the re-orientation of this lipid product to the other membrane leaflet occurs in the ER and secretory pathway via the actions of a scramblase and P4-ATPase flippase^24^.

Beyond revealing the mechanism of MBOAT1’s catalysis, the structures provided here may facilitate the development of specific inhibitors of MBOAT1. Since PS remodeling appears crucial for modulating ferroptosis^4^ and apoptosis^25,26^, such inhibitors may provide valuable tools to probe MBOAT1-dependent lipid states and may have therapeutic potential in diseases linked to MBOAT1-dependent ferroptosis, such as estrogen receptor–positive breast cancers^4^, or altered PS remodeling, such as endometriosis^27^.

## Acknowledgements

We thank members of the Farese and Walther laboratory, especially Dr. Xiaojun Xiang, for helpful discussions. We thank M. Jason de la Cruz and Dr. Sagnik Sen at the MSKCC Structural Biology Core Facility for assistance with cryo-EM data acquisition, and Dr. Cem Komurcuoglu at the MSKCC High-Performance Computing Center for computational support for cryo-EM data processing. We also thank Dr. Rui Yan, Dr. Nicholas Spellmon and the team at the Howard Hughes Medical Institute Janelia Cryo-EM Facility for cryo-EM microscope operation and data collection. We thank Gary Howard for editorial assistance.

## Author contributions

L.K., R.V.F. and T.C.W. conceived the project. L.K. expressed and purified proteins, prepared cryo-EM grids, collected and processed cryo-EM data, built atomic models from single-particle cryo-EM maps, prepared samples for lipidomic analysis, and performed enzymatic activity assays. S.K. performed and analyzed MD simulations. R.L. performed lipidomic experiments and analyzed the lipidomics data. L.C.A., P.L., and X.J. assisted with cellular experiments. D.K. advised on cryo-EM data processing and project development. L.K. and T.C.W. wrote the manuscript with input from all authors.

## Methods

### Protein expression and purification

cDNA with the sequence of full-length human MBOAT1 (UniProt: Q6ZNC8) was synthesized and cloned into a modified pEG-BacMam vector containing a C-terminal 3C protease cleavage site and followed by a FLAG tag. H381A and N350A mutations were introduced by site-directed mutagenesis. All proteins were expressed in HEK293S GnTI^-s^uspension cells (ATCC) by baculovirus-mediated transduction. Cells were cultured in FreeStyle 293 Expression Medium (Gibco, 12338026) supplemented with 2% (v/v) FBS and maintained at 37 °C with 8% CO_2_. Recombinant baculoviruses were generated and amplified using the Bac-to-Bac Baculovirus Expression System protocol. For protein expression, HEK293S GnTI^-c^ells at a density of 2.0–3.0 million cells mL^-1^ were transduced with 4.0–4.5% (v/v) P3 baculovirus. After 18 h, 10 mM sodium butyrate was added, and the culture temperature was lowered to 30 °C. At 48 h after sodium butyrate addition, cells were harvested by centrifugation at 550 g. Cell pellets were resuspended in lysis buffer containing 20 mM Tris-HCl pH 8.0, 150 mM NaCl, 10% (v/v) glycerol, supplemented with cOmplete EDTA-free protease inhibitor cocktail (Roche, 04693132001) and 1 mM phenylmethylsulfonyl fluoride. Cells were disrupted by sonication, and membranes were solubilized by addition of 2% (w/v) glycol-diosgenin (GDN; Anatrace, GDN101) for 1–2 h at 4°C with gentle agitation. Insoluble material was removed by centrifugation at 33,745 g for 1 h at 4 °C. The clarified supernatant was incubated with anti-FLAG M2 resin (Sigma-Aldrich, A2220) for 1 h at 4 °C. The resin was then packed in a gravity-flow column (Bio-Rad, 7321010) and washed with 10 column volumes (CV) of buffer A (20 mM Tris-HCl pH 8.0, 150 mM NaCl, 0.02% (v/v) GDN). Then, MBOAT1 protein was eluted with 5 CV of elution buffer (20 mM Tris pH8.0, 150 mM NaCl, 0.02% (w/v) GDN, 0.150 mg mL^-1^ 3xFLAG peptide). The eluted protein was concentrated and further purified by size-exclusion chromatography (SEC) on a Superose 6 Increase column 10/300 (Cytiva Life Science, 29-0915-96) equilibrated in buffer A.

### Cryo-EM sample preparation and data collection

Freshly purified WT MBOAT1 was concentrated to 7.45 mg mL^-1^ and applied directly to cryo-EM grids without addition of exogenous substrate. For substrate-bound mutant samples, MBOAT1 H381A at 3.2 mg mL^-1^ was incubated with 167 μM 16:0 lyso-PS (Avanti Polar Lipids, 858142P-1mg) for 20 min before vitrification. MBOAT1 N350A at 2.5 mg mL^-1^ was incubated with 140 μM 16:0 lyso-PS and 140 μM oleoyl coenzyme A (Sigma-Aldrich, O1012-10MG) for 20 min before vitrification. For the PS-bound sample, oleoyl-CoA was added to the WT MBOAT1 to a final concentration of 250 μM and incubated on ice for 30 min before vitrification. The final protein concentration was 5.32 mg mL^-1^.

Cryo-EM grids were prepared by applying 3.5 μL of protein sample to freshly glow-discharged Quantifoil R1.2/1.3, 400-mesh gold holey carbon grids (Quantifoil, Q4100AR1.3). Grids were blotted and plunge-frozen in liquid ethane using a Vitrobot Mark IV (Thermo Fisher Scientific) operated at 4°C and 100% humidity. Vitrified grids were stored in liquid nitrogen until data acquisition.

Cryo-EM data for WT MBOAT1 were collected on a Titan Krios electron microscope (Thermo Fisher Scientific) operating at 300 keV at the HHMI Janelia Research Campus. Movies were recorded using a K3 detector (Gatan) equipped with a BioQuantum energy filter (Gatan) operated with a 20-eV slit width. Data were collected in super-resolution mode using SerialEM at a nominal magnification of 105,000x, corresponding to a physical pixel size of 0.827Å. The total electron exposure was approximately 60 e^-Å-2,^ and the defocus range was -0.6 to -1.7 μm.

Cryo-EM data for MBOAT1 H381A, MBOAT1 N350A and PS-bound MBOAT1 were collected on a Titan Krios electron microscope (Thermo Fisher Scientific) operated at 300 keV at the Richard Rifkind Center for Cryo-EM, Memorial Sloan Kettering Cancer Center. Movies were recorded in counting mode on a Falcon 4i direct electron detector (Thermo Fisher Scientific) equipped with a Selectris X energy filter (Thermo Fisher Scientific), using automated data collection in Smart EPU. Images were collected at a nominal magnification of 165,000x, corresponding to a physical pixel size of 0.725Å. The total dose exposure was approximately 60 e^-Å-2^. The defocus range was -0.7 to -1.8 μm. Data collection parameters are summarized in Table 1.

### Electron microscopy data processing

Cryo-EM data processing was performed in cryoSPARC software (v4.5.0-v4.7.0)^28^. Movies were gain-normalized and motion-corrected using patch-motion-correction algorithm. Contrast transfer function (CTF) parameters were estimated using Patch CTF estimation, and micrographs were manually curated on the basis of CTF quality and estimated resolution. Data processing workflows are summarized in Extended Fig. 2.

Briefly, particles were initially picked from a subset of micrographs using blob-based auto-picking and subjected to several rounds of two-dimensional (2D) classification. Selected particles were then used to train Topaz models for neural network-based particle-picking^29^. False-positive particles and poorly defined classes were excluded through iterative rounds of 2D classification, *ab-initio* reconstruction, and heterogenous refinement. Particle subsets showing well-defined transmembrane features were selected for non-uniform refinement. Because of the small size of MBOAT1, detergent micelle density was subtracted to improve alignment and map quality. A tight mask excluding the detergent micelle was generated and used for signal subtraction, followed by local refinement to obtain the final reconstructions.

Local resolution estimations, Fourier shell correlation (FSC) validations using the gold-standard FSC criterion (0.143 and 0.5) and viewing direction distribution analysis were calculated in CryoSPARC. Comprehensive data processing and statistics are summarized in Table 1.

### Model building and refinement

An AlphaFold 3 model of human MBOAT1 was used as the initial model for building into the cryo-EM density maps^30^. Subsequently, the models were manually rebuilt using Coot to accurately fit into experimental densities^31^. Atomic coordinates were refined against the corresponding cryo-EM maps using PHENIX real-space refinement with secondary-structure, geometry and Ramachandran restraints^32^. Ligand geometry restraints were generated from canonical SMILES strings using eLBOW in PHENIX^33^.

Model-to-map agreement was assessed by FSC analysis between cryo-EM maps and corresponding final atomic models. Model validation was performed using MolProbity within PHENIX^34^. A detailed model refinement and validation statistics are summarized in Table 1. Structural figures were prepared using UCSF Chimera, UCSF ChimeraX, and PyMOL (Schrödinger)^35,36^.

### Cell culture

T47D-Cas9 and T47D-gMBOAT1 cells were cultured in RPMI medium (Gibco, 61870127) supplemented with 10% (v/v) FBS, 100 U mL^-1^ penicillin/streptomycin (Gibco, 15140122), and maintained at 37 °C with 5% CO_2_. Cells were harvested by scraping in 2 mL of ice-cold PBS, collected by centrifugation at 500 g, and snap-frozen in liquid nitrogen. The cell pellets were suspended in ice-cold PBS and lysed using a microtip ultrasonic homogenizer. Protein concentrations were determined using a BCA assay, and sample amounts were normalized according to protein concentration during extraction and subsequent LC-MS analysis. The T47D cell lines were a gift from the laboratory of Xuejun Jiang.

### Lipid profiling of protein-associated lipids

Chromatographic separation was performed using a Vanquish Horizon UHPLC system equipped with an Accucore C18 column (2.1 × 150 mm, 2.6 µm; Thermo Fisher Scientific) coupled to a Thermo Scientific Orbitrap Exploris 240 mass spectrometer. The column compartment was maintained at 50 °C and the autosampler at 15 °C. Mobile phase A consisted of acetonitrile/water (60:40, v:v) and mobile phase B of isopropanol/acetonitrile/water (88:10:2, v/v/v); both phases contained 10 mM ammonium formate and 0.1% formic acid. The flow rate was held constant at 0.3 mL/min, and the injection volume was 5 µL. A linear gradient of mobile phase B was applied as follows: 10% B at 0 min, increasing to 86% B at 20 min, and further increasing to 95% B at 26 min, where it was held until 29 min. The gradient was then returned to initial conditions (10% B) at 29.1 min and held for 3 min to re-equilibrate the column, for a total run time of 32 min.

Full-scan MS¹ spectra were acquired at 120,000 resolution over a scan range of *m/z* 200– 1,700, with a maximum injection time of 50 ms and standard AGC target. Data-dependent MS² scans (ddMS²) were acquired at 30,000 resolution using a cycle time of 1.5 s, with a maximum injection time of 54 ms, isolation width of 1.0 *m/z*, intensity threshold of 20,000, and dynamic exclusion of 3 s (5 ppm mass tolerance), in both positive and negative mode for each sample. Source parameters were as follows: spray voltage +3,500 V (positive mode) and −2,800 V (negative mode), RF lens 25%, sheath gas 25, auxiliary gas 10, and sweep gas 1 (arbitrary units), vaporizer temperature 200 °C, and ion transfer tube temperature 250 °C.

### Untargeted lipid profiling of WT and KD cells

Chromatographic separation was performed using a Vanquish Horizon UHPLC system (Thermo Fisher Scientific, Waltham, MA) equipped with an Accucore C30 column (2.1 × 150 mm, 2.6 µm; Thermo Fisher Scientific) coupled to a Thermo Scientific Orbitrap ID-X Tribrid mass spectrometer. The column compartment was maintained at 45 °C and the autosampler at 15 °C. Mobile phase A consisted of acetonitrile/water (60:40, v/v) and mobile phase B of isopropanol/acetonitrile/water (88:10:2, v/v/v); both phases contained 10 mM ammonium formate and 0.1% formic acid. The flow rate was held constant at 0.25 mL/min, and the injection volume was 3 µL. A linear gradient of mobile phase B was applied as follows: 30% B from −3 to 0 min (column equilibration), increasing to 43% B at 2 min, 55% B at 2.1 min, 65% B at 12 min, and 85% B at 18 min, before reaching 100% B at 20 min, where it was held until 25 min. The gradient was then returned to initial conditions (30% B) at 25.1 min and held until 28 min to re-equilibrate the column, for a total run time of 28 min.

Full-scan MS¹ spectra were acquired in polarity-switching mode at 120,000 resolution over a scan range of *m/z* 200–1,700, with a maximum injection time of 50 ms and standard AGC target. Data-dependent MS² scans were acquired at 30,000 resolution using an AcquireX iterative identification workflow for lipid annotation, with a maximum injection time of 54 ms, isolation width of 1.0 *m/z*, intensity threshold of 20,000, and dynamic exclusion of 3 s (5 ppm mass tolerance). Source parameters were as follows: spray voltage +3,500 V (positive mode) and −3,000 V (negative mode), RF lens 50%, sheath gas 50, auxiliary gas 10, and sweep gas 2 (arbitrary units), vaporizer temperature 300 °C, and ion transfer tube temperature 320 °C.

### Data processing

Raw data files were processed using LipidSearch 5.2 (Thermo Fisher Scientific, Waltham, MA) for lipid identification and annotation, with peak integration and quantitation performed in Skyline (MacCoss Lab, University of Washington)^37^. LipidCruncher^38^ was used for downstream filtering and quality control of annotated lipid features and for figure generation, with additional statistical analysis and visualization performed in MetaboAnalyst^39^.

### Fluorescence-based catalytic assay

Purified MBOAT1 acyltransferase activity was measured using a fluorescence-based assay that detects free CoA-SH released from acyl-CoA during the reaction. Free CoA-SH was detected using 7-diethylamino-3-(4-maleimidophenyl)-4-methylcoumarin (CPM).

Reactions were performed in a final volume of 20 μL containing 2 mg mL⁻¹ BSA, 0.6 μM purified MBOAT1, 100 μM oleoyl-CoA and 100 μM 16:0 lysophospholipid. Reactions were incubated at 37 °C for 20 min for endpoint assay. For Michaelis–Menten analysis, oleoyl-CoA was maintained at 100 μM and the concentration of 16:0 lysophospholipid was varied. Reactions used for kinetic analysis were incubated at 37 °C for 3 min, within the linear range of product formation. Reactions were quenched by addition of SDS to a final concentration of 1.3% and incubated at room temperature for 5 min. 180 μL of CPM working solution (50 μM in 20 mM Tris pH 8.0, and 150 mM NaCl) was added to the reaction, and the mixture was kept at room temperature for 15 min and followed by fluorescence detection with excitation at 355 nm and emission at 460 nm. Background signals from control reactions lacking enzyme, lysophospholipid or oleoyl-CoA were subtracted. Nonlinear regression to the Michaelis-Menten equation and allosteric sigmoidal fitting were performed using GraphPad Prism 10.

### Molecular dynamics simulation

The resolved MBOAT1–PS complex was embedded in a lipid bilayer, with each leaflet consisting of 100 POPC lipids, using^40^ mstool. The system was solvated and neutralized with 0.15 M NaCl, energy-minimized, and equilibrated prior to production. Simulations were performed with the Martini 3 coarse-grained force field^41^ in GROMACS 2024.6 using a 20-fs integration timestep. For MBOAT1, an elastic network with a force constant of 700 kJ mol⁻¹ nm⁻² and an upper cutoff of 0.9 nm was applied to preserve the tertiary structure. Pressure was maintained at 1 bar with the Parrinello-Rahman barostat^42^ using semi-isotropic coupling, a compressibility of 4.5 × 10⁻⁵ bar⁻¹, and a 12 ps time constant. Temperature was maintained at 310 K with the V-rescale thermostat^43,44^ using a 1-ps time constant. A relative dielectric constant of 15 was used, and Coulomb interactions were modeled with a reaction field, with a nonbonded cutoff of 1.1 nm. A simulation was run for 800 ns each.

Trajectories were analyzed with MDAnalysis^45^ and visualized with ChimeraX^36^. Occupancy was calculated from the per-frame distance between every bead of each protein residue and every PS bead. A protein residue and a PS bead were considered in contact if any inter-bead distance fell below a cutoff of 0.5 nm. Contacts were scored as binary per frame and accumulated over the 800 ns analyzed (800 frames), then normalized by the number of analyzed frames to yield an occupancy between 0 and 1, defined as the fraction of frames in which the contact was present.

### Statistics and reproducibility

Statistical analyses were performed using GraphPad Prism 10. Data were presented as mean ± standard deviation. The number of biological replicates for *in vitro* and *in vivo* experiments are indicated in the figure legends.

## Data and code availability

All atomic models and cryo-EM maps have been deposited in the Protein Data Bank (PDB) and in the Electron Microscopy Data Bank (EMDB): MBOAT1 WT (PDB 37FF, EMD-78139); MBOAT1 H381A (PDB 37FG, EMD-78140); MBOAT1 N350A (PDB 37FH, EMD-78141); PS-bound MBOAT1 (PDB 37FI, EMD-78142).

---

**Extended Data Fig. 1.**
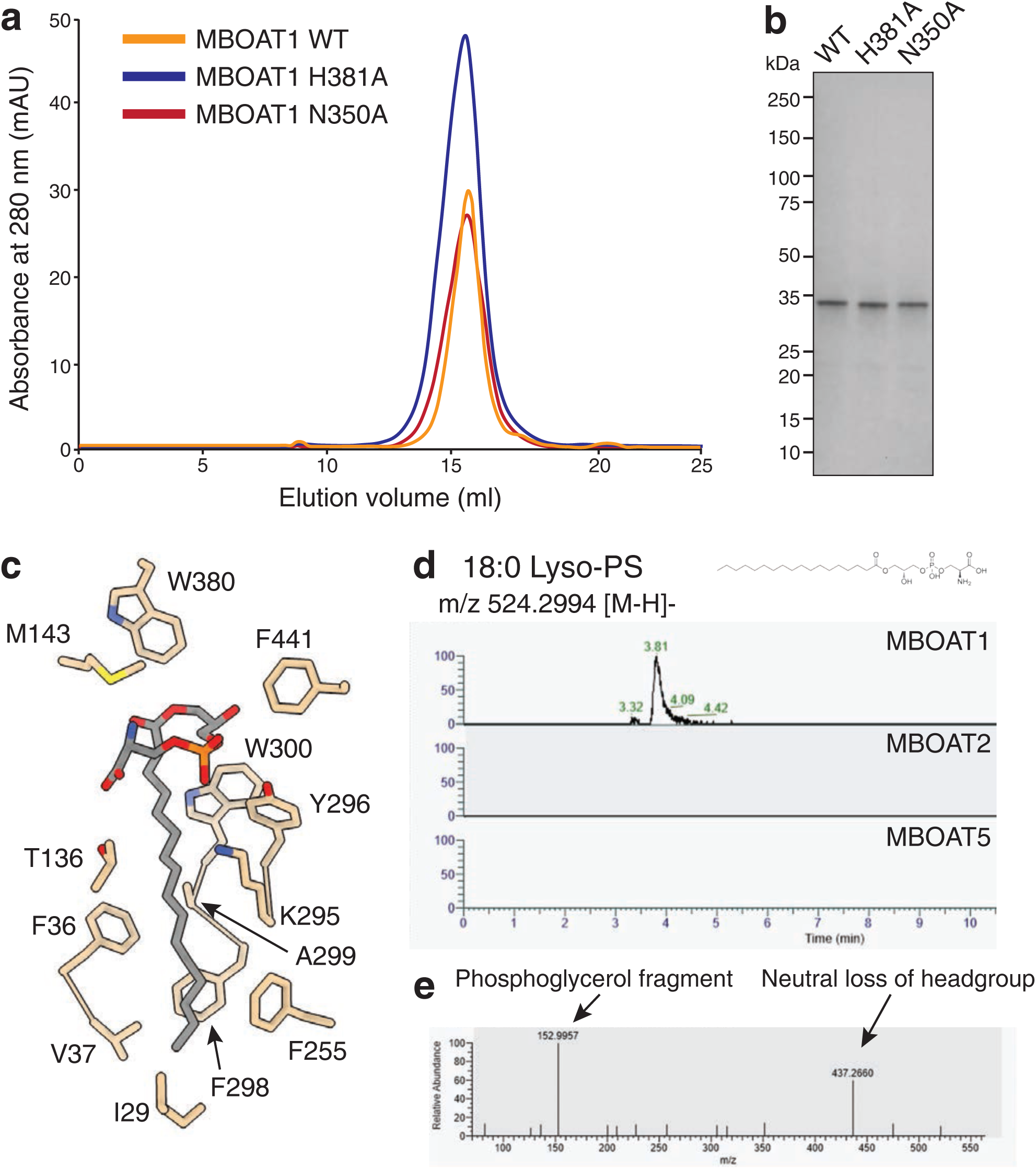
Biochemical characterization of purified MBOAT1 and associated endogenous lipid. **a,** Representative size-exclusion chromatography profiles of human MBOAT1 WT, H381A, and N350A. The three proteins showed similar monodisperse elution profiles. **b,** SDS-PAGE analysis of the purified MBOAT1 WT, H381A and N350A peak fractions. **c,** Interactions between lyso-phospholipid acyl chain and MBOAT1 residues. The non-proteinaceous density was modelled as an endogenous lyso-PS, and surrounding residues are shown as sticks. **d,** LC-MS chromatograms demonstrating signal detected for 18:0 lyso-PS (negative mode) in purified MBOAT1 and absence in MBOAT2 and MBOAT5. **e,** MS2 spectra of 18:0 lyso-PS with characteristic structural fragments from headgroup loss and phosphoglycerol fragment.

**Extended Data Fig. 2.**
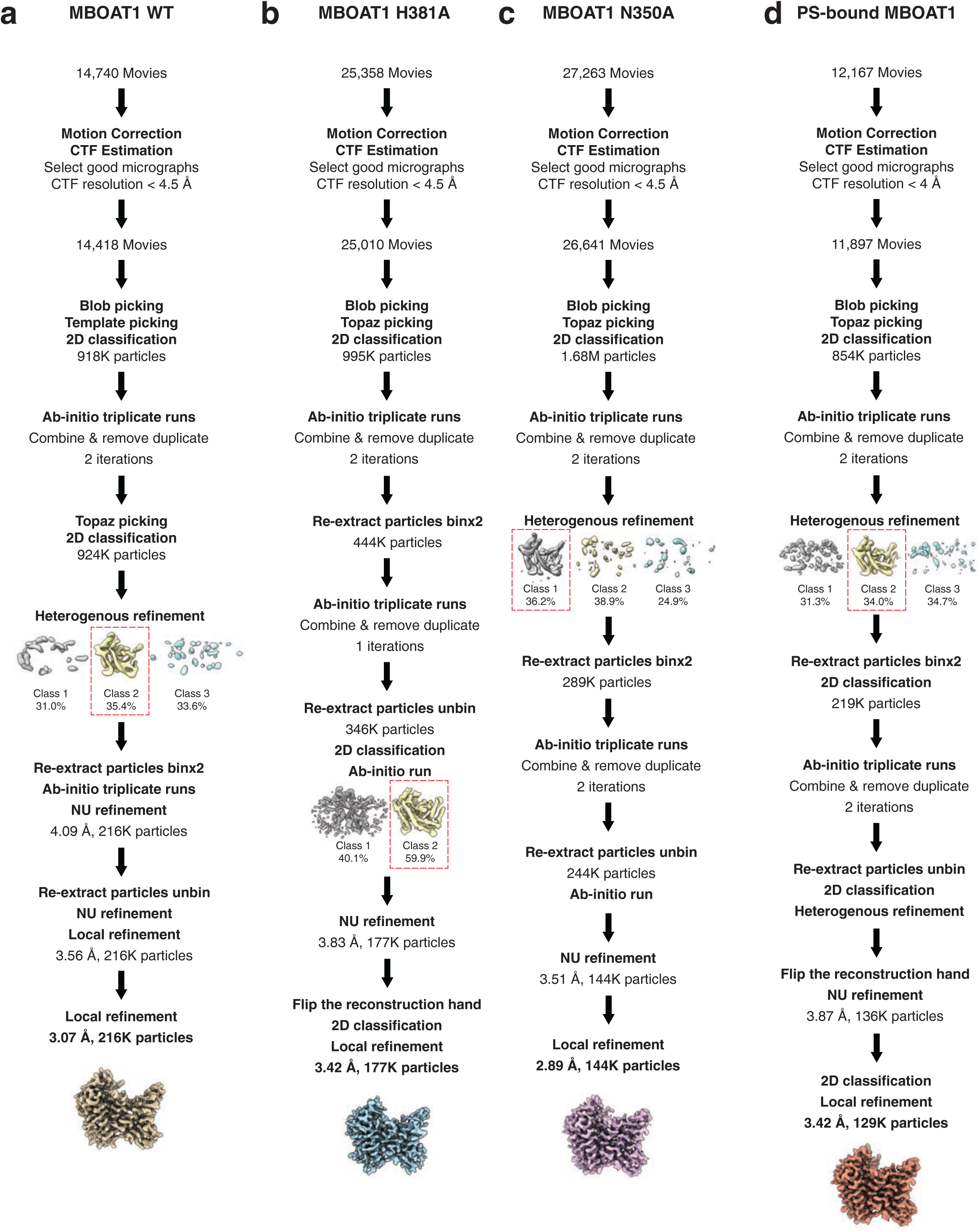
Cryo-EM data processing workflow. **a-d,** Overview of the cryo-EM image processing pipeline employed to determine MBOAT1 structures presented in this study. Detailed procedures are described in the Methods.

**Extended Data Fig. 3.**
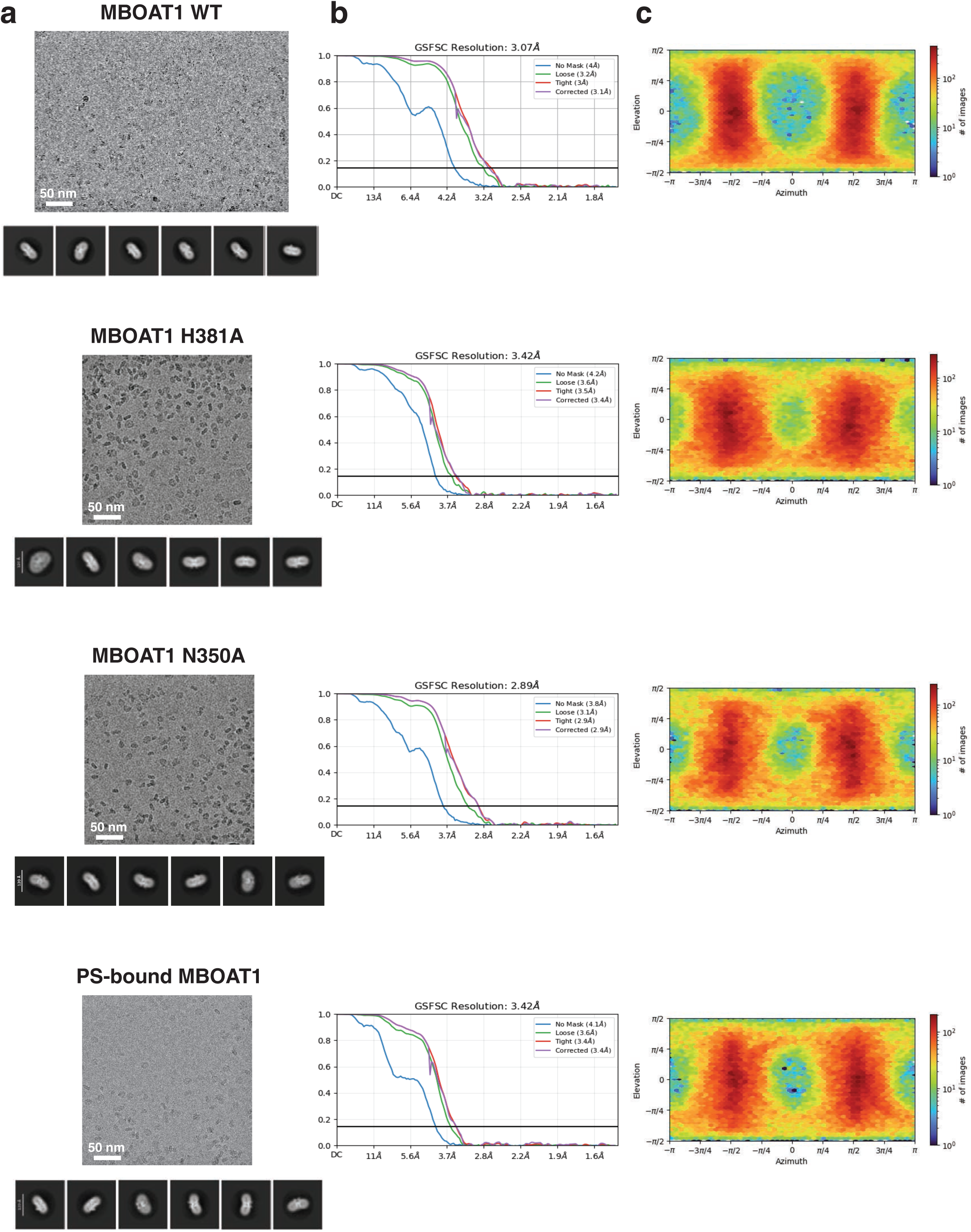
Representative micrographs, 2D class averages, and FSC curves. **a,** A representative micrograph, six 2D class averages are presented for each dataset. **b,** Fourier shell correlation (FSC) plots of the final map are presented, respectively. FSC curves between two half-maps were calculated using the 0.143 criterion. **c,** The distribution of particle orientations of the final map.

**Extended Data Fig. 4.**
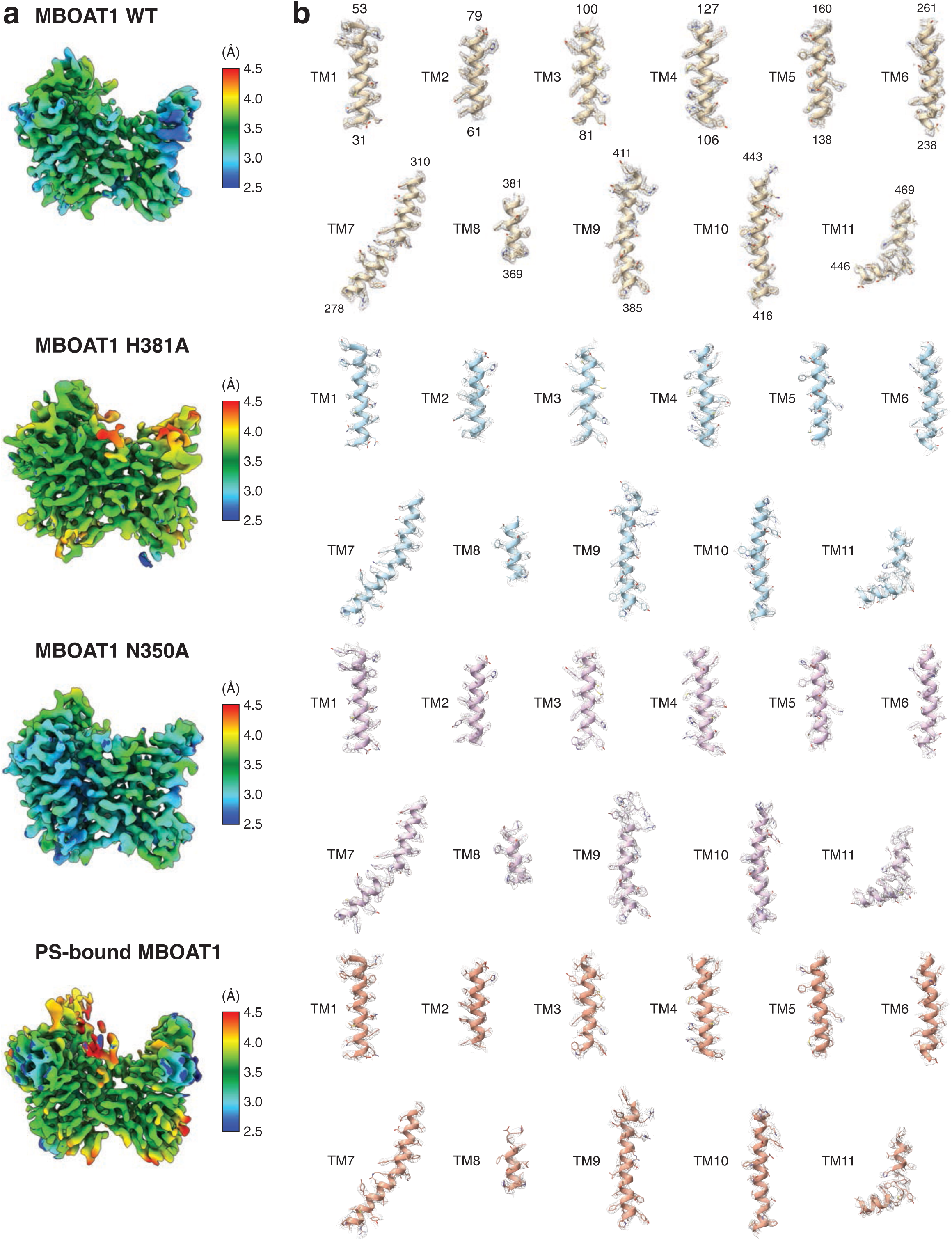
Quality of cryo-EM map. **a,** Final cryo-EM maps colored according to local resolution estimates. The color scale indicates local resolution in Å, as shown in the accompanying bar. **b,** Densities corresponding to the transmembrane domain (TMD) regions of each structure are displayed as grey meshes overlaid with the corresponding atomic models.

**Extended Data Fig. 5.**
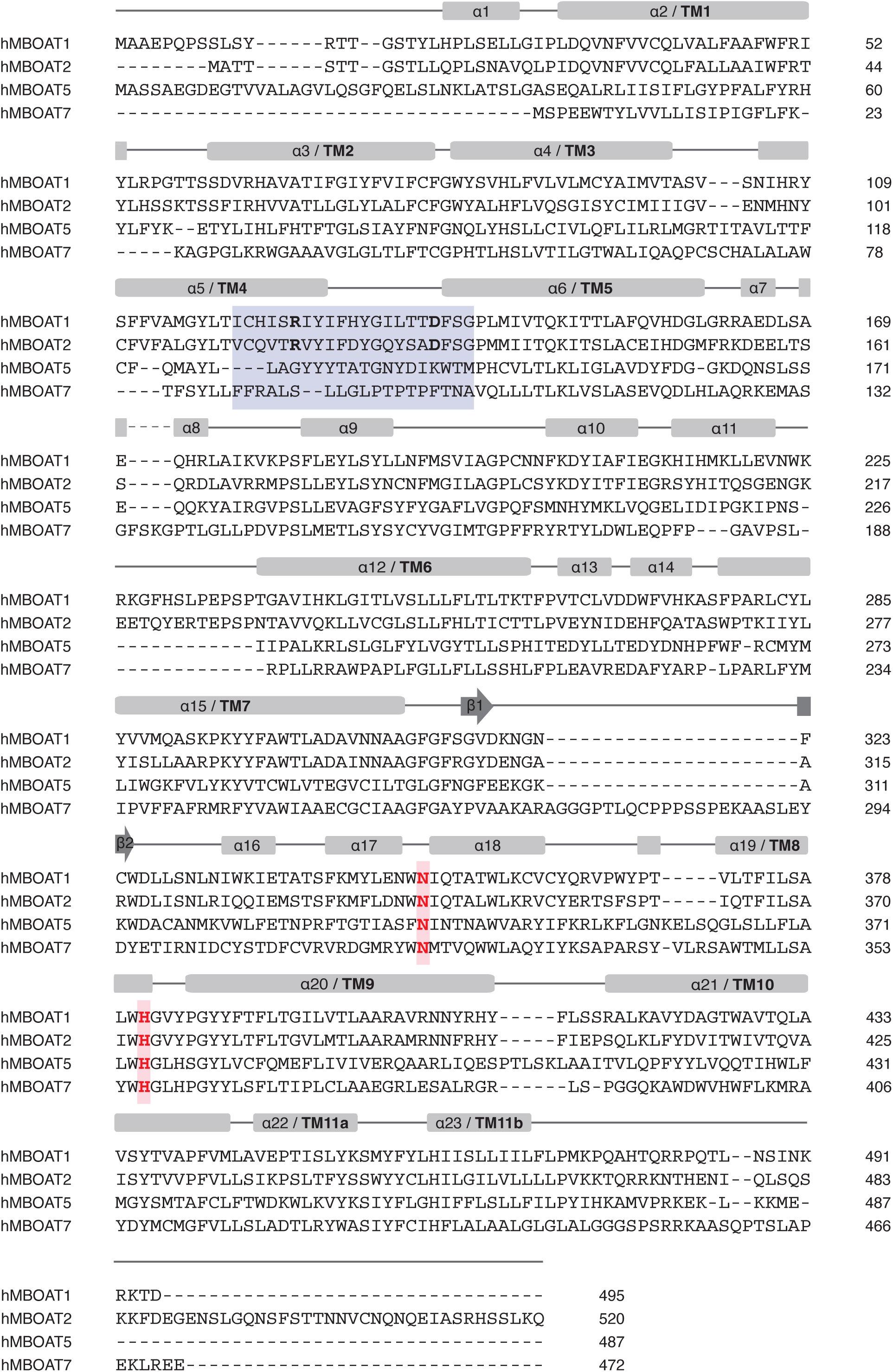
Sequence alignment of human MBOAT proteins. Sequence alignment of hMBOAT1, hMBOAT2, hMBOAT5, and hMBOAT7. Secondary-structure elements of hMBOAT1 are shown above the alignment, with transmembrane helices labelled according to the hMBOAT1 structure. The TM4–TM5 selectivity loop region, corresponding to residues 120–140 in hMBOAT1, is highlighted in blue. Conserved catalytic residues, including Asn350 and His381 in hMBOAT1, are highlighted in red.

**Extended Data Fig. 6.**
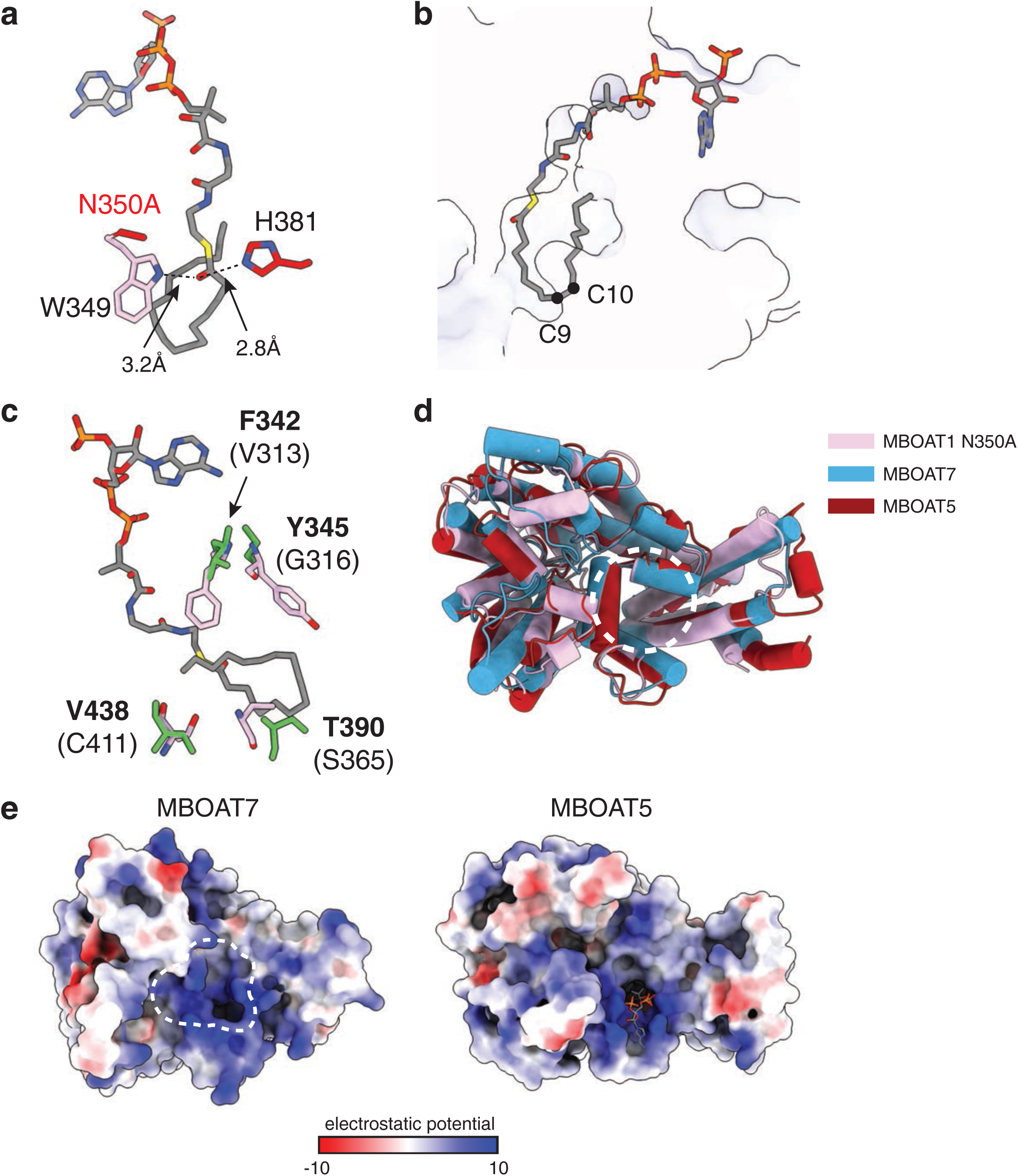
Comparison of acyl donor-binding tunnel among MBOAT proteins. **a,** Close-up view of oleoyl-CoA bound to MBOAT1 N350A. The thioester positions adjacent to the catalytic residues Asn350 and His381. Distances between the thioester-proximal region and catalytic site residues are indicated in dash [[“indicated by dashed lines]]. **b,** View of the MBOAT1 acyl donor-binding tunnel occupied by oleoyl-CoA. The oleoyl chain extends into the membrane-embedded tunnel. The C9-C10 cis double bond of the oleoyl chain is indicated as dots. **c,** Comparison of the MBOAT1 oleoyl-CoA binding tunnel with the corresponding region of MBOAT7. Selected MBOAT1 residues that narrow the donor acyl-binding tunnel are shown in pink, and the corresponding MBOAT7 residues are shown in green and indicated in parentheses. **d,** Structural superposition of MBOAT1 N350A, MBOAT7 and MBOAT5 (PDB: 7F40). MBOAT1 N350A: pink, MBOAT7: blue and MBOAT5: red. **e,** Electrostatic surface potentials of MBOAT7 and MBOAT5 at the cytoplasm-facing acyl donor-binding tunnel. Dashed outlines indicate the putative CoA-binding entrance of MBOAT7. Electrostatic potentials are colored from −10 kTe^−1^ in red to +10 kTe^−1^ in blue.

**Extended Data Fig. 7.**
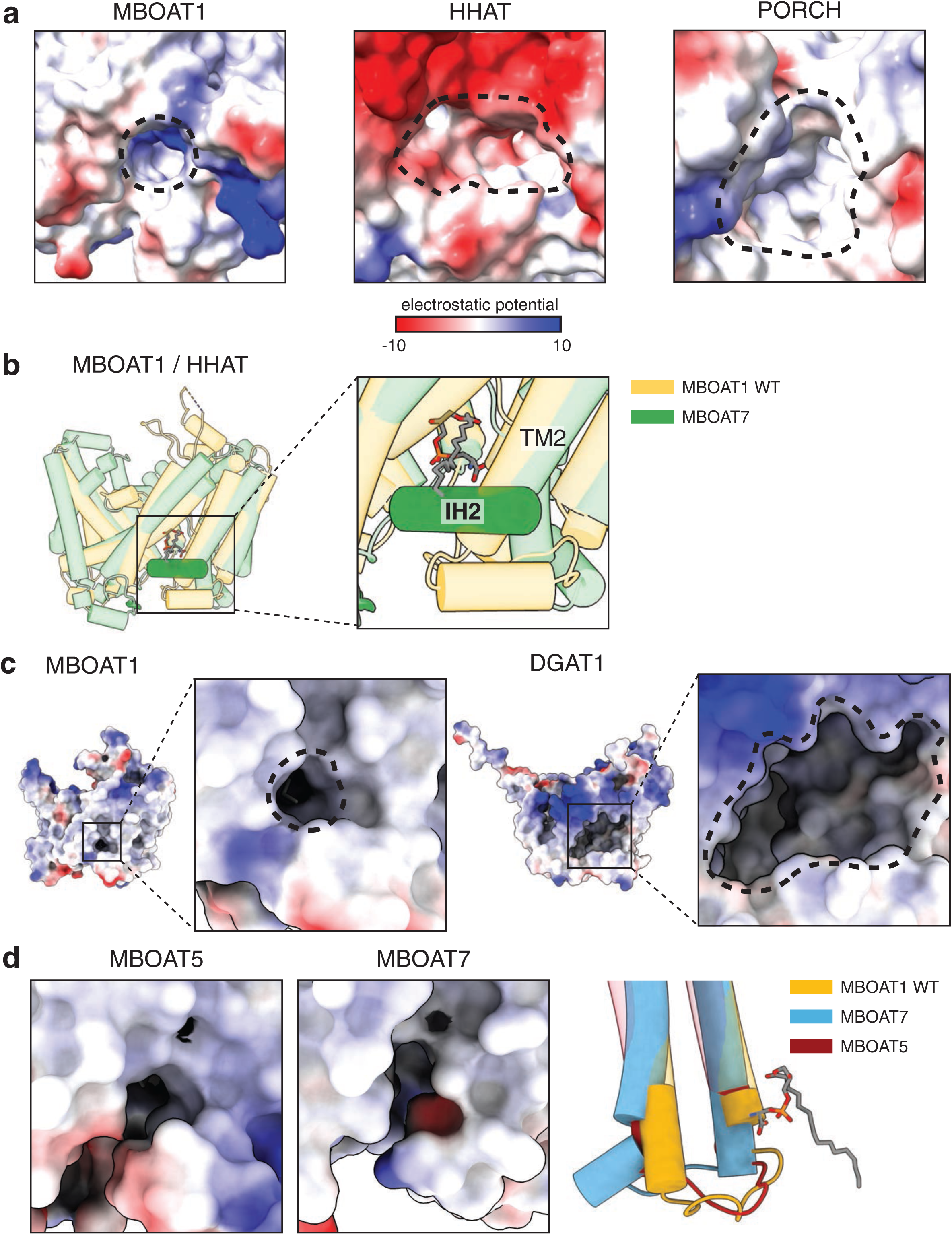
Divergent substrate access pathways among MBOAT family enzymes. **a,** Electrostatic surface potentials of MBOAT1, HHAT and PORCN viewed from the lumen-facing side. Putative lumenal openings are indicated by dashed outlines. Electrostatic potentials are colored from −10 kTe^−1^ in red to +10 kTe^−1^ in blue. **b,** Structural comparison of MBOAT1 and HHAT around the lateral-access region. MBOAT1 is shown in yellow and HHAT in green. The TM2-linked IH2 helix of HHAT occupies the region corresponding to the MBOAT1 lateral gate. **c,** Electrostatic surface potentials of MBOAT1 and DGAT1 viewed from the membrane-facing lateral side. Dashed outlines indicate the lateral openings. DGAT1 contains a broader lateral opening than MBOAT1, consistent with accommodation of a bulky diacylglycerol acceptor. **d,** Comparison of the lateral gate and selectivity loop regions of lipid-remodeling MBOAT enzymes. Left, electrostatic surface potentials of MBOAT5 and MBOAT7 around the corresponding lateral gate. Right, structural comparison of the selectivity loop regions of MBOAT1, MBOAT5 and MBOAT7. MBOAT1 is shown in yellow, MBOAT7 in blue and MBOAT5 in red.

**Extended Table 1.** Cryo-EM data collection, refinement and validation statistics.

|  | MBOAT1 WT<br>(EMDB-78139)<br>(PDB 37FF) | MBOAT1 H381A<br>(EMDB-78140)<br>(PDB 37FG) | MBOAT1 N350A<br>(EMDB-78141)<br>(PDB 37FH) | POPS-bound<br>MBOAT1<br>(EMDB-78142)<br>(PDB 37FI) |
| --- | --- | --- | --- | --- |
| **Data collection and processing** |  |  |  |  |
| Magnification | 105,000 | 165,000 | 165,000 | 165,000 |
| Voltage (kV) | 300 | 300 | 300 | 300 |
| Electron exposure (e-/Å<sup>2</sup>) | 60 | 60 | 60 | 60 |
| Defocus range (μm) | -0.6 to -1.7 | -0.7 to -1.8 | -0.7 to -1.8 | -0.7 to -1.8 |
| Pixel size (Å) | 0.827 | 0.725 | 0.725 | 0.725 |
| Symmetry imposed | C1 | C1 | C1 | C1 |
| Initial particle images (no.) | 924,000 | 995,000 | 1,680,000 | 854,000 |
| Final particle images (no.) | 216,000 | 177,000 | 144,000 | 129,000 |
| Map resolution (Å) | 3.07 | 3.42 | 2.89 | 3.42 |
| FSC threshold (0.143) |  |  |  |  |
| **Refinement** |  |  |  |  |
| Initial model used (PDB code) | De novo | 37FF | 37FF | 37FF |
| Model resolution (Å) | 3.5 | 3.7 | 3.1 | 4.0 |
| FSC threshold (0.5) |  |  |  |  |
| Map sharpening *B* factor (Å<sup>2</sup>) | -49.6 | -131.5 | -83.7 | -95.9 |
| Model composition |  |  |  |  |
| Non-hydrogen atoms | 3,701 | 3,653 | 3,719 | 3,780 |
| Protein residues | 453 | 450 | 449 | 460 |
| Ligands | LKS: 1 | LKS: 1 | LKS: 1<br>3VV: 1 | POP: 1 |
| *B* factors (Å<sup>2</sup>) |  |  |  |  |
| Protein | 68.43 | 81.03 | 45.55 | 60.06 |
| Ligand | 86.32 | 83.26 | 58.67 | 71.37 |
| R.m.s. deviations |  |  |  |  |
| Bond lengths (Å) | 0.003 | 0.003 | 0.004 | 0.002 |
| Bond angles (°) | 0.540 | 0.545 | 0.607 | 0.540 |
| Validation |  |  |  |  |
| MolProbity score | 1.72 | 2.19 | 2.11 | 1.91 |
| Clashscore | 5.82 | 6.30 | 4.58 | 5.45 |
| Poor rotamers (%) | 0.51 | 4.62 | 3.08 | 2.24 |
| Ramachandran plot |  |  |  |  |
| Favored (%) | 93.99 | 95.06 | 91.46 | 94.98 |
| Allowed (%) | 6.01 | 4.49 | 8.54 | 5.02 |
| Disallowed (%) | 0.00 | 0.45 | 0.00 | 0.00 |

